# Sea lamprey nests promote the diversity of benthic macroinvertebrate assemblages

**DOI:** 10.1101/2022.09.04.506553

**Authors:** Marius Dhamelincourt, Jacques Rives, Marie Pons, Aitor Larrañaga, Cédric Tentelier, Arturo Elosegi

## Abstract

The habitat heterogeneity hypothesis states that increased habitat heterogeneity promotes species diversity through increased availability of ecological niches. We aimed at describing the local-scale effects of nests of the sea lamprey (*Petromyzon marinus* L.) as ecosystem engineer on macroinvertebrate assemblages. We hypothesized that increased streambed physical heterogeneity caused by sea lamprey spawning would modify invertebrate assemblages and specific biologic traits and promote reach-scale diversity. We sampled thirty lamprey nests of the Nive River in three zones: the unmodified riverbed (control) and zones corresponding to the nest: the area excavated (pit) and the downstream accumulation of pebbles and cobbles (mound). The increased habitat heterogeneity created by lamprey was traduced by biological heterogeneity with a reduced density of invertebrates (1160 to 6540 individuals per *m*^2^ in control, 680 to 6460 individuals per *m*^2^ in pit and 1980 to 6240 individuals per *m*^2^ in mound) and number of taxa (23.5 ± 3.9 taxa for control, 18.6 ± 3.9 taxa in pit and 21.2 ± 4.5 taxa for mound) in the pit compared to other zones. However the overall taxa diversity in nest increased with 82 ± 14 taxa compared to the 69 ± 8 taxa estimated in control zone. Diversity indices were consistent with the previous results indicating a loss of *α* diversity in pit but a higher *β* diversity between a pit and a mound than between two control zones, especially considering Morisita index accounting for taxa abundance. Trait analysis showed high functional diversity within zones with a reduced proportion of collectors, scrapers, shredders, litter/mud preference and small invertebrates in mound, while the proportion of “slabs, blocks, stones and pebbles” preference and largest invertebrates increased. Pit presented the opposite trend, while control had globally intermediate trait proportions. Our results highlight important effects on species and functional diversity due to habitat heterogeneity created by a nest-building species, what can ultimately influence food webs and nutrient processes in river ecosystems.

## Introduction

Habitat heterogeneity has been widely recognized as an important factor structuring biological assemblages [1, 2]. The habitat heterogeneity hypothesis [3] states that increased habitat heterogeneity promotes species diversity thanks to a greater number of available ecological niches. Li and Reynolds [4] defined heterogeneity in an operational way as “the complexity and/or variability of a system property in space and/or time”. Heterogeneity can be considered at different scales, and depending on the scale considered, it has contrasted effects on ecological processes [5]. In the case of stream invertebrates, substrate heterogeneity can affect food distribution [6] and thus condition their local-scale (i.e. nest-scale) feeding and movements [7], whereas movements at larger scales can respond to drift if there are no available interstices or substrate for attachment.

In aquatic ecosystems, heterogeneity analyzes are often used to explain macroinvertebrate assemblages. Beisel et al. [8] found that the number of taxa in 0.5 to 4 m radius zones was higher in a heterogeneous environment with diverse substrate and an elevated patchiness. Furthermore, the same study found that homogeneous environments are more prone to the dominance of few taxa, probably due to the lack of competition with taxa from neighbouring patches. At the sample scale, heterogeneous substrate hosts more diverse assemblages as found by Boyero [9] for 225 cm^2^ samples. Substrate appears to be a key driver of macroinvertebrate assemblage structure followed by current velocity and depth [10]. However, increased heterogeneity does not necessarily enhance macroinvertebrate diversity, at least in stream restoration projects [11].

Factors influencing heterogeneity include abiotic and biotic processes, such as the interaction between hydraulics and the substrate, or the effects of engineer species [12]. Together, they affect habitat heterogeneity, which in turn can affect ecosystem functioning [6]. Substrate disturbance by foraging fish is another cause of substrate heterogeneity, which may destabilize the substrate and cause an increased bedload transport during subsequent flood events [13].

Fish nesting, especially in species such as salmonids, which move large volumes of sediment and alter local bed morphology, is a biotic process that can increase riverbed substrate heterogeneity [14]. For migratory species with high nest density in the spawning grounds, the ecological effects can be important and cascade to affect biological assemblages and ecosystem processes. For instance, Moore and Schindler [15, 16] found that bed disturbance by salmon in spawning grounds caused severe seasonal declines in periphyton and benthic invertebrate abundance, with no recovery within the same season. However, other species such as chubs of the genus *Nocomis* build nests that promote macroinvertebrate density and even allow some taxa to persist in degraded streams [17]. Migratory species likely to affect substrate heterogeneity and the assemblages include the sea lamprey (*Petromyzon marinus* L.), which builds 40-220 cm-wide nests [18, 19, 20] in late spring [21]. Both male and female lamprey remove cobbles with their mouth and release them downstream so that the resulting nest consists of a pit carpeted with sand or gravel, followed by a mound of pebbles and cobbles. This structural heterogeneity might have functional consequences, as the density of some taxa such as Hydropsychidae, Philopotamidae and Heptageniidae can increase up to tenfolds in mounds [22] in an oligotrophic stream. Another study [23] found an increased Simuliidae density in areas with disturbed substrate and added sea lamprey carcasses. These results suggest an effect of sea lamprey nesting activity on macroinvertebrate assemblages, with potential functional consequences.

Here, we aimed at describing the local-scale effects of sea lamprey as ecosystem engineer on macroinvertebrate assemblages at the nest. We expect the taxonomic and functional diversity on the nest as a whole (pit+mound) to be higher than on undisturbed substrate. We hypothesized that increased streambed physical heterogeneity caused by sea lamprey nesting would promote invertebrate diversity and specific biologic traits. Pits, with deeper and more lentic conditions than unmodified riverbed (control) would attract lentophilic and fine sediment fauna, whereas the shallower and more lotic mounds would attract taxa with opposite preferences. In addition, nests with more heterogeneous current and depth may host more diverse communities. To test our hypotheses, we first assessed the variability of depth and current velocity on pit, mound and control as these variables are among the most likely to explain differences in macroinvertebrate assemblages [10]. Then, we compared the density, taxa richness and diversity of macroinvertebrates across nest zones. Finally, we studied ecological traits describing food type, mode of alimentation, substrate preference and body size, to infer the effects of nests on invertebrate functional heterogeneity.

## Methods

### Study site and experimental period

The study took place in the Nive River, a 79 km long river situated in Northern Basque Country, France, and draining a basin of 1030 km^2^. The selected section corresponds to a reach of 400 m long, located in Saint-Martin-d’Arrossa (43° 14’ 34.926” N, 1° 18’ 27.305” W) and bypassed by a hydropower scheme. It is mainly composed of riffles and runs suitable for sea lamprey nesting, as shown by the 46 nests found on a previous survey [24]. Macroinvertebrate samples were collected on July 7 2020, at the end of the reproductive season, to avoid disturbing sea lamprey spawning. No individuals were observed in the two weeks prior to sampling, a period of colonization short enough to limit the risk of flooding but sufficient to allow for stability of local invertebrate assemblages [25].

### Macroinvertebrate samples and nest characteristics

We studied 30 completed nests where no lamprey was observed. For each nest, we defined three zones: the **control zone**, 20 to 50 cm upstream from the upstream verge of the nest pit, indicated the initial conditions of the habitat chosen by the lamprey to build their nest. The second zone was the **pit**, delimited as the area excavated by the lamprey, where water was deeper and the substrate finer than on the control. The third zone was the **mound**, corresponding to the downstream accumulation of pebbles and cobbles. For all three zones current velocity (± 1 cm/s) was measured at 5 cm from the bottom (using a magnetic flowmeter “FLO-MATE 2000”) and water depth (± 1 cm) was measured with a Vernier gauge. Transversal (perpendicular to the current) and longitudinal (parallel to the current) diameters of the pit were measured using a measuring tape (± 1 cm).

Macroinvertebrates were sampled using a Surber net (1/20 *m*^2^, 0,5 mm mesh) by stirring the entire top layer of the sediment. Three samples, one per zone, were collected on each nest. After collection the samples were stored in 70° ethanol up to the determination. Most invertebrates were identified following AFNOR standard XP T 90-388 at the genus level except for Oligochaeta (subclass), Diptera (family) and some individuals for whom genus determination was not certain (determined at the family level). Macroinvertebrate traits were determined following Tachet et al. [26] and expressed as the frequency of individuals belonging to a certain trait over the total number of individuals in the sample. We selected food type, mode of alimentation, substrate preference and body size as these traits are likely to vary according to the physical characteristics of the nests. Food type and mode of alimentation directly reflect the resources provided by a given environment. With different current and substrate characteristics those traits should vary between zones. As substrate will differ, substrate preferences may reflect the sorting made by spawners between zones, if the scale of the nest has an impact on macroinvertebrate spatial location. Finally, body size emphasizes the carrying capacity for smaller or bigger invertebrates, which completes the substrate preference.

### Invertebrate diversity

We used several *α* and *β* indices to measure the effects of lamprey nests on the diversity of macroinvertebrate assemblages. For α indices we calculated the indices in each zone to compare the pit and the mound with the control. First, we measured the species richness with the specpool function of the vegan package [27, version 2.5-7] using a Chao1 Eq [28]. Then, Shannon α diversity index [29, Eq 1] indicated how diverse the taxa in each zone are, increasing with the increase of evenness and taxa richness in a sample. Shannon index was completed with Pielou index [30, Eq 2] to specifically analyze the evenness of the assemblages. These indices were transformed into Log Response Ratio (LRR), corresponding to the effect size of the diversity differences between nest (pit/mound) zones and control zones (Eq 3 and Fig 4) where the “index modified zone” corresponds either to the index of a pit or to the index of a mound. Indices were calculated for each pit and each mound and compared to the mean of Shannon and Pielou indices obtained for the control zones (“index control zone”). Behind this analysis, we compared how a given index (Shannon or Pielou) evolved on a pit or on a mound - thus reflecting a disturbed habitat - compared to the average value of these indices in the section considered.

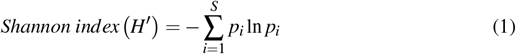

**Fig 1.**
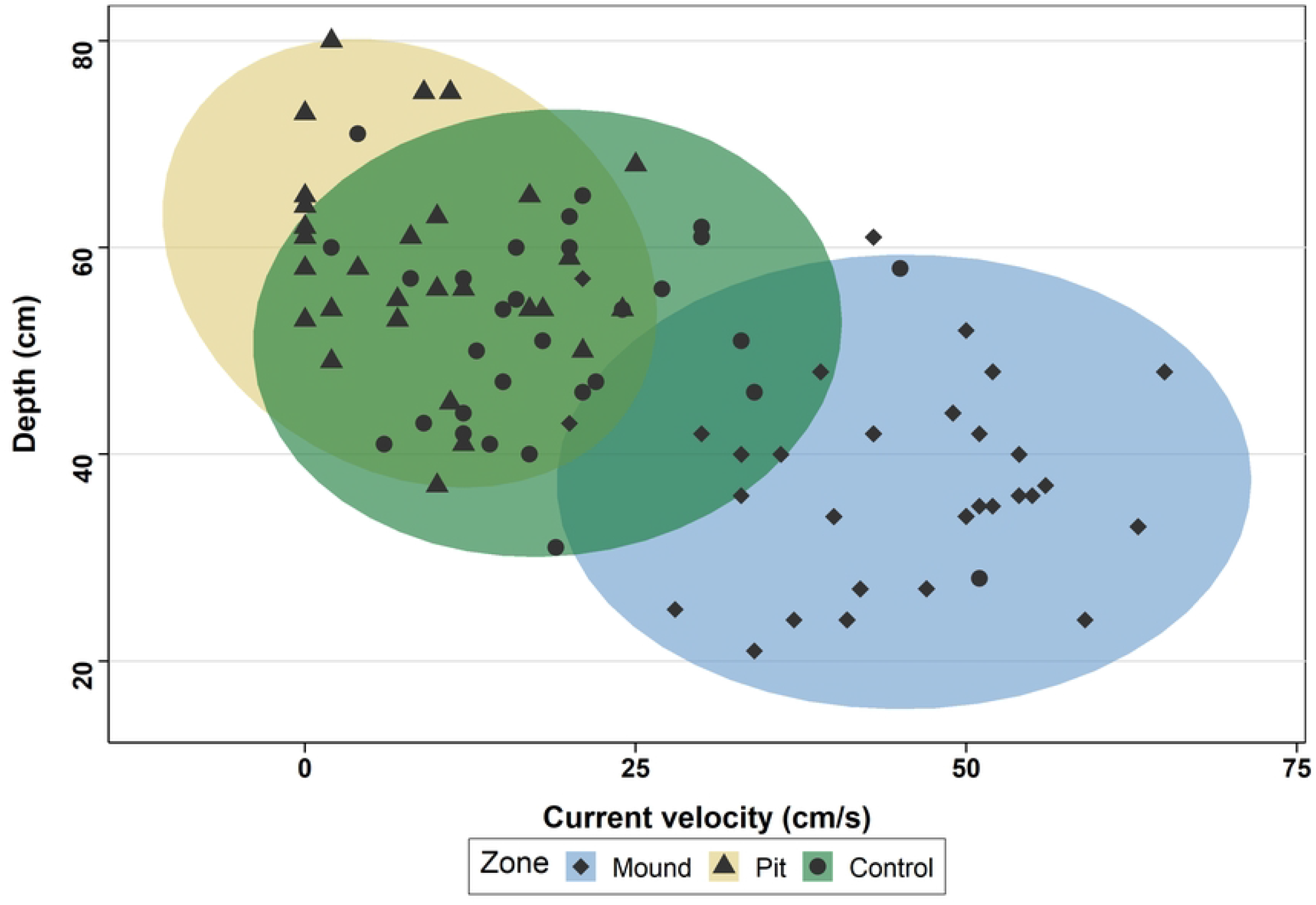
Relationship between depth and current velocity for each nest at each zone. Ellipses correspond to multivariate t-distribution.

**Fig 2.**
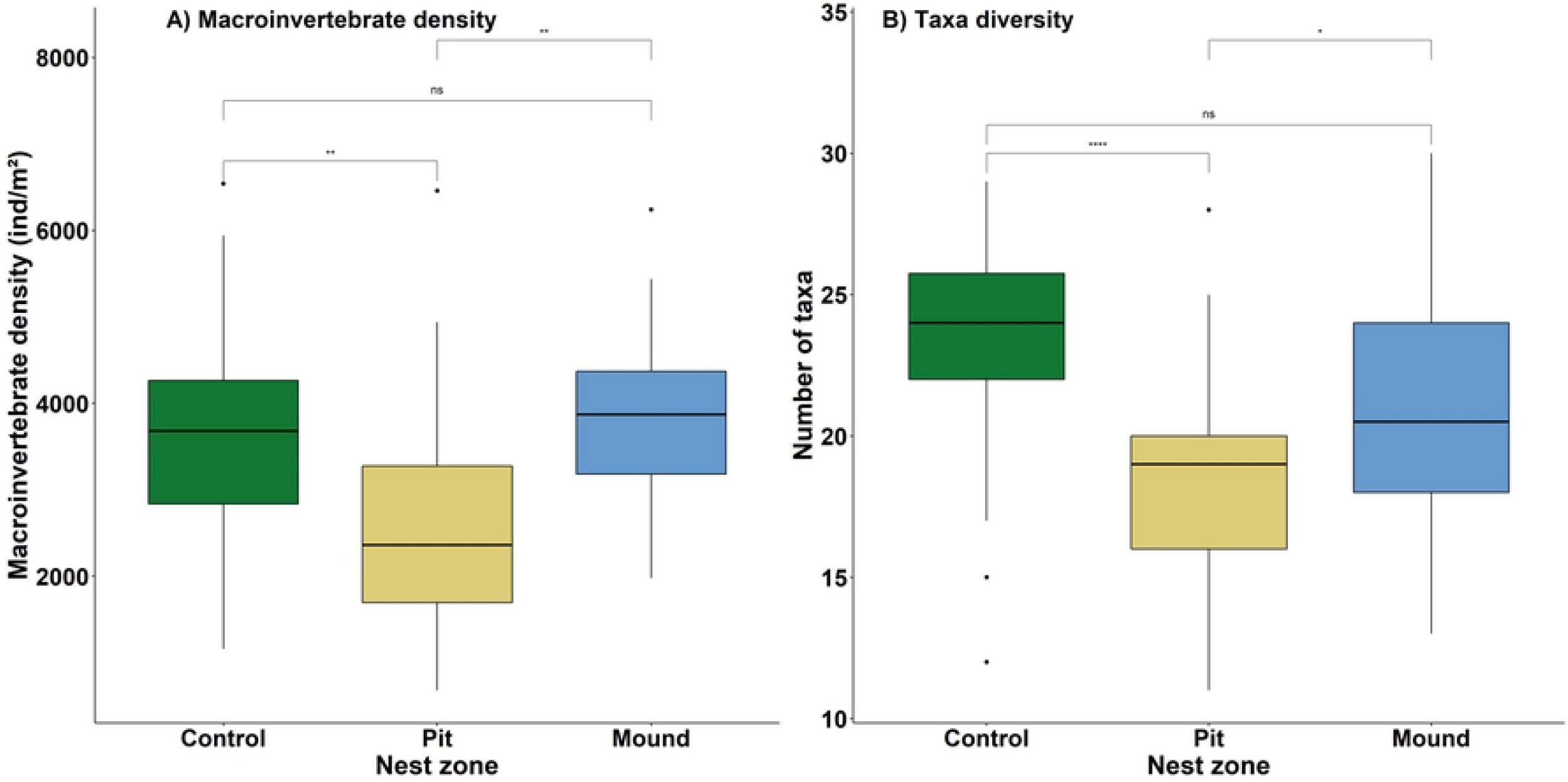
Density of macroinvertebrates (*ind/m*^2^) (A) and number of taxa per sample (B) for the three zones studied. With *ns P* > 0.05, \**P* ≤ 0.05, \*\**P* ≤ 0.01, \*\*\*\**P* ≤ 0.0001.

**Fig 3.**
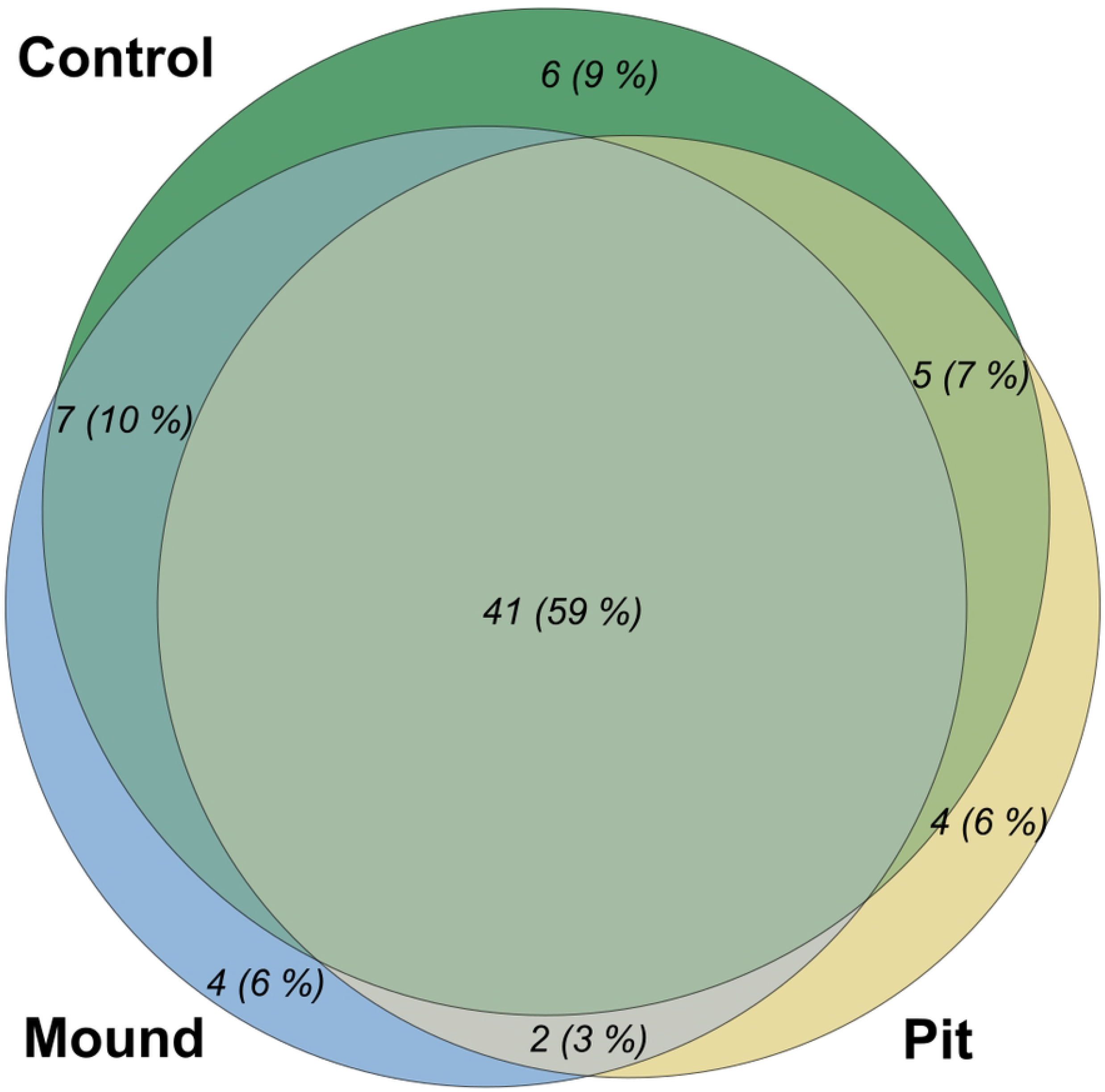
Venn diagram with the distribution of taxa among zones. The area of each part is proportional to the number of taxa indicated in absolute number and percentage of total taxa richness.

**Fig 4.**
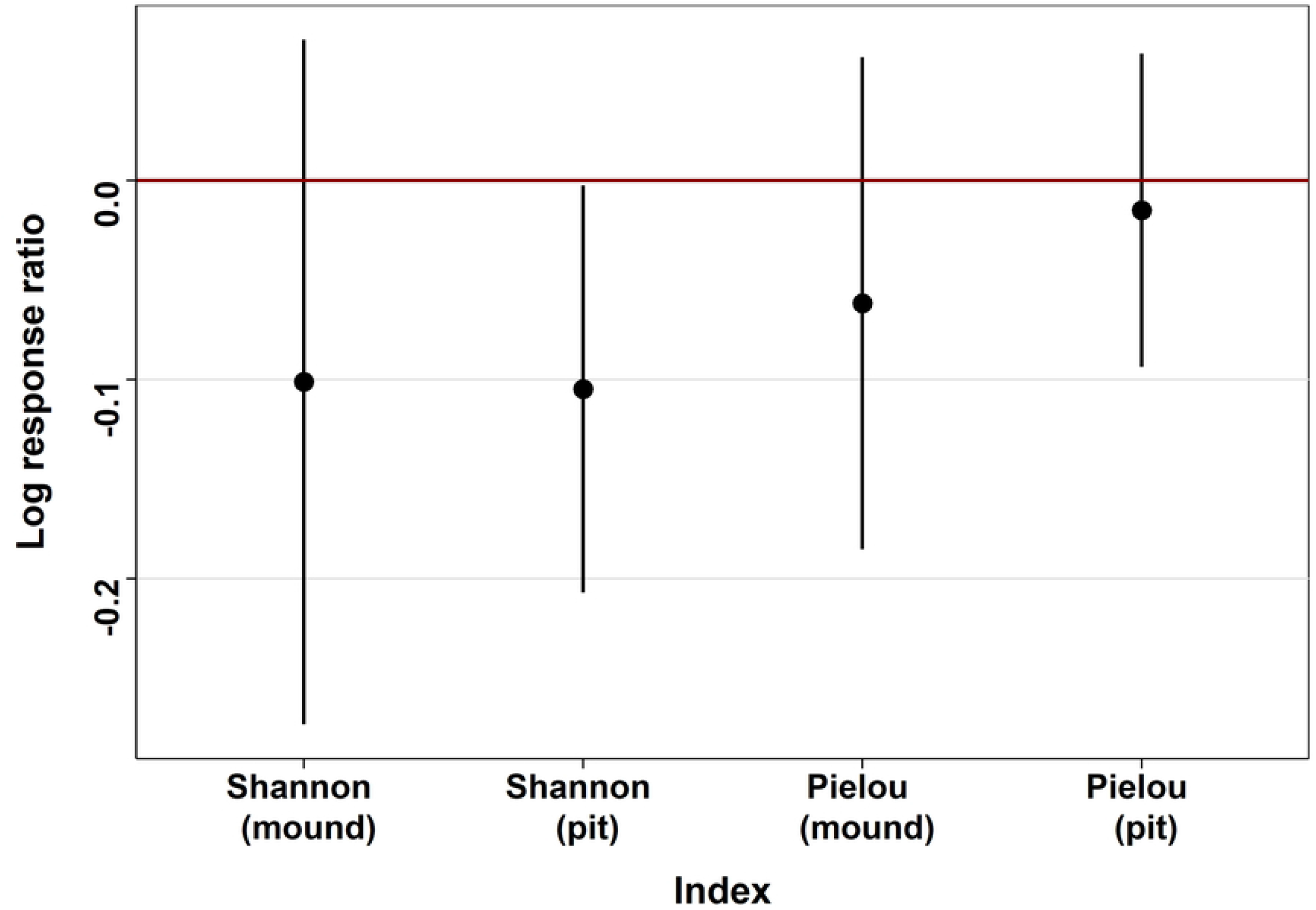
Log response ratios of Shannon and Pielou *α* diversity indices for each pit and each mound compared to the mean of Shannon and Pielou indices obtained for the control zones. Dots correspond to the mean and error bars correspond to the standard deviation.

*Where *p_i_* is the proportion of a taxon i on the total number of individuals N and S is the total number of taxa in the sample*.

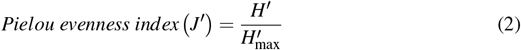

*Where H’is the Shannon diversity index and* 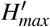 *is the maximum possible value of H’ (if every species was equally likely)*: 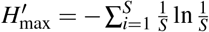 *and S being the total number of taxa in the sample*.

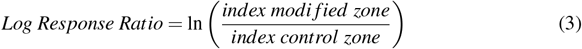

Following this *α* diversity overview we studied several *β* diversity aspects to compare nest and control. For each index we calculated all the pairwise differences between pit and mound and between two control zones. Firstly, we calculated Sorensen dissimilarity index [31, Eq 4] which includes both replacement and nestedness and corresponds to a global indicator of taxa diversity. Then we calculated Simpson *β* diversity index to study taxa replacement [31, Eq 5].

The second component of the Sorensen index, the taxa nestedness, was obtained by computing the difference between Sorensen and Simpson indices [32]. Finally, to test the diversity considering taxa abundance and not solely their occurrence, we calculated the Morisita dissimilarity index [33, Eq 6]. As for the *α* diversity indices the *β* indices were transformed into Log Response Ratio (Eq 3 and Fig 5) where the “index modified zone” corresponds to a pairwise index between a pit and a mound and the “index control zone” to a pairwise index between two controls. This LRR indicates if the diversity difference between two locations of a nest (always a pit versus a mound) differs from the same comparison made between two controls, thus reflecting a difference in taxa heterogeneity. To avoid a nest effect we did not compare the taxa of a pit versus the mound of the same nest and therefore we randomly selected the pit and the mound to be compared in each index value.

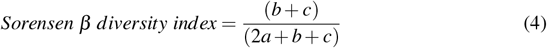

**Fig 5.**
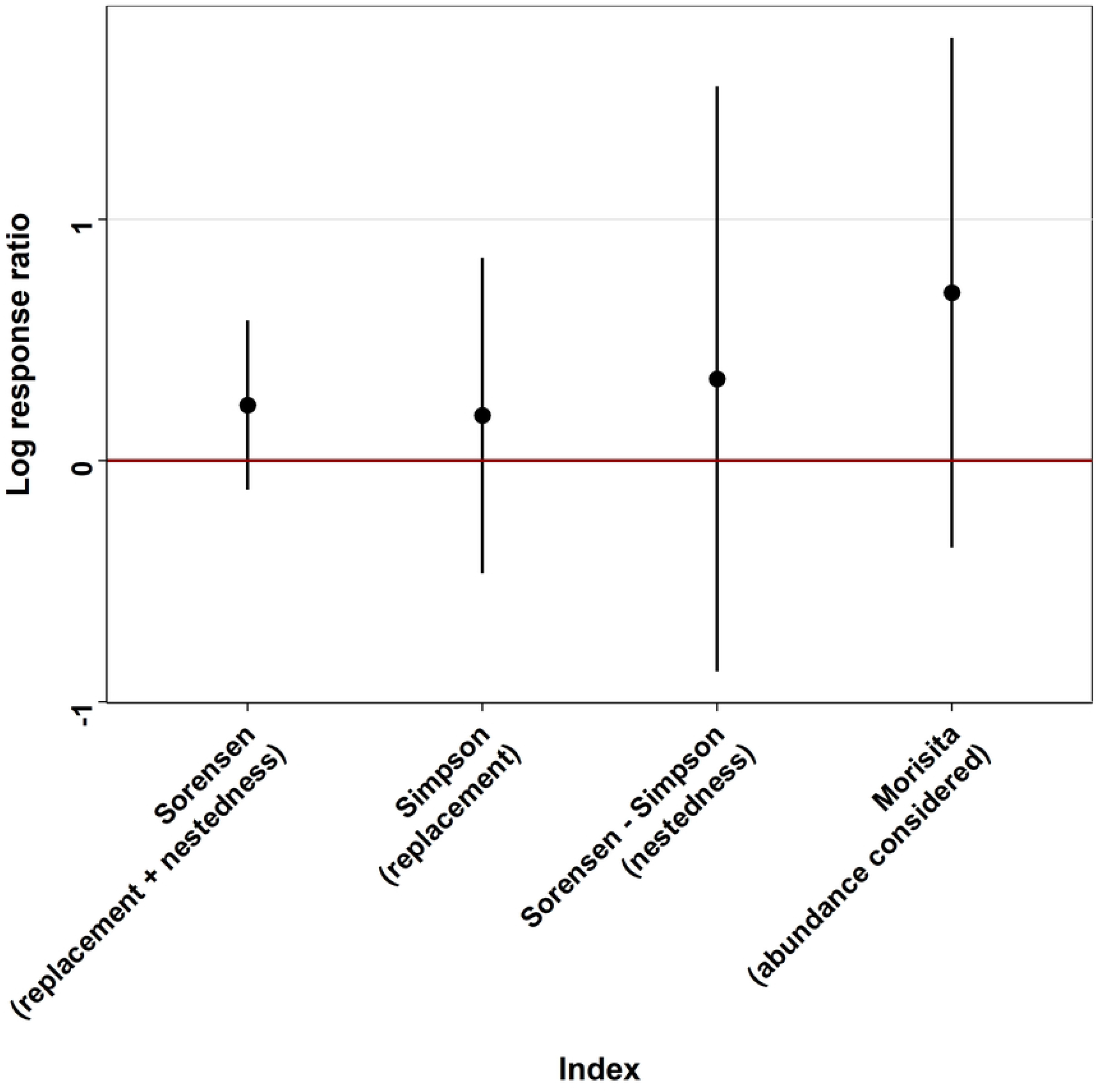
Log response ratios of *β* diversity indices of all mound and pit pairwise indices, each compared to a pairwise control index among all possible control pairwise comparisons (control zone randomly selected, not directly upstream from the pit and mound considered). Dots correspond to the mean and error bars correspond to the standard deviation.

*Where a is the number of common taxa between two sites, b the number of taxa exclusive to a site, and c the number of taxa exclusive to the other site*.

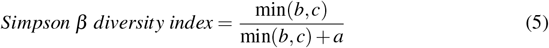

*Where min (b, c) is the lowest number of exclusive taxa between two sites, b being the number of taxa exclusive to a site, and c the number of taxa exclusive to the other site; a is the number of common taxa between the two sites*.

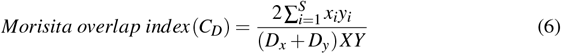

*Where x_i_, is the number of times taxon i is represented in the total X from one site; y, is the number of times taxon i is represented in the total Y from another site; D_x_ and D_y_ are the Simpson’s α diversity index for the x and y sites respectively*:

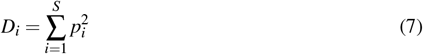

*With S the total number of taxa in the sample and p_i_, the proportion of a taxon i on the total number of individuals N*.

### Data analysis

All analyzes were performed using R software version 4.1.0 [34] and a significance level of 0.05.

The significance of differences between nest zones for depth, current, density, diversity, Shannon index and Pielou index was determined using linear mixed model and the *lmer* function of the *lme4* package [35] with a random effect of nest identity. For density and taxa diversity (Fig 2A and B) a pairwise Tukey test was used to check the differences between each zone rather than globally. Trait differences between zones were determined using binomial mixed models with control or mound as reference condition and nest identity as a random effect (function *glmer* of the *lme4* package; Bates et al. [35]. The links between trait and nest characteristics were studied using Redundancy analysis (RDA), a multivariate approach of linear models applied in *vegan* package [27, version 2.5-7] with *rda* function and setting traits as response variables. Traits were implemented after Hellinger transformation to give low weight to traits with low counts and many zeros. Explanatory variables were the five variables describing nest characteristics: transversal and longitudinal diameter, depth, current and depth difference between pit and mound. The latter was calculated assuming nests with higher depth difference could be more heterogeneous with a more important volume and habitat complexity. To test the effect of current and depth on abundance and taxa diversity, a general linear model was used with current and depth as covariates and following a Gaussian distribution.

## Results

### Nests characteristics

Lamprey nests differed greatly in their dimensions, transversal diameter of the pit averaging 129.2 cm (range, 70-210 cm) and longitudinal diameter 117,4 cm (range, 50-260 cm, Table 1).

**Table 1.** Summary of the nests characteristics measured in the study; mean ± sd (cv).

|  | Pit | Mound | Control |
| --- | --- | --- | --- |
| Transversal diameter (cm) | $129.2 \pm 37.2$ (0.3) | - | - |
| Longitudinal diameter (cm) | $117.4 \pm 45.7$ (0.4) | - | - |
| Depth (cm) | $59.2 \pm 9.9$ (0.2) | $38.3 \pm 10.1$ (0.3) | $51.8 \pm 10.1$ (0.2) |
| Current (cm/s) | $8.8 \pm 7.8$ (0.9) | $43.8 \pm 11.6$ (0.3) | $19.8 \pm 11.1$ (0.6) |
| Depth difference (cm) | $20.9 \pm 4.8$ (0.2) | | - |

The pit was the deepest zone (59.2 ± 9.9 cm) followed by the control (51.8 ± 10.1 cm) and the mound (38.3 ± 10.1 cm), and differences were statistically significant considering mixed models p-values (Table 2). The coefficient of variation for depth was similar for all three zones (Table 2). Current velocity ranged from 0 to 65 cm/s (Table 1) and varied among zones inversely to depth (Fig 1), although the coefficient of variation was larger. Again, differences were statistically significant considering mixed models results.

**Table 2.** Mixed model results for analyzes of differences between zones for nest characteristics, density and diversity of macroinvertebrates and *α* diversity indices, with nest identity as a random effect. With \**P* ≤ 0.05, \*\**P* ≤ 0.01, \*\*\**P* ≤ 0.001.

| Variables | *Df* | $\chi^2$ | *P* |
| --- | --- | --- | --- |
| Depth | 2 | 503.5 | <2e-16 *** |
| Current | 2 | 278.41 | <2e-16 *** |
| Density | 2 | 17.475 | 1.61e-4 *** |
| Diversity | 2 | 25.736 | 2.58e-6 *** |
| Shannon index (control/pit/mound) | 2 | 12.739 | 1.71e-3 ** |
| Pielou index (control/pit/mound) | 2 | 6.6937 | 3.52e-2 * |

### Macroinvertebrate density and diversity

The density of macroinvertebrates (Fig 2A) ranged from 680 to 6460 individuals per m^2^ in pit, from 1980 to 6240 individuals per *m*^2^ in mound and from 1160 to 6540 individuals per *m*^2^ in control. It was significantly lower in pit than in control and mound, but there were no differences between control and mound. The taxa richness (Fig 2B) also did not vary significantly between control (23.5 ± 3.9 taxa) and mound (21.2 ± 4.5 taxa) but was higher in these two zones than in pit (18.6 ± 3.9 taxa). The Venn diagram (Fig 3), describing exclusivity or sharing of taxa, indicates an important taxa overlap with 59 % of the taxa found on all zones. The significant pattern of reduced taxa richness found at the nest scale was not found by summing all the different taxa identified in the thirty samples of each zone and visualizing it in the Venn diagram.

### Diversity

The estimate of Chao1 diversity index was 82 ± 14 taxa for nest (pit + mound) and 69 ± 8 taxa for the control zone. In spite of the slight overlap of the standard error of those estimates the species richness seems to be higher in the nest than in the control. Log response ratios of the *α* diversity indices (Fig 4) showed an overall trend of reduced diversity and equitability for both Shannon and Pielou indices in pit and mound, compared to the average values observed in the control zone. However, only the log response ratio of the Shannon index of the pit is strictly below 0 and indicates a significantly reduced diversity in pit compared to the control zone. The log response ratios of *β* diversity indices (Fig 5) showed a higher overall *β* diversity between a pit and a mound than between two control zones. However the log response ratios were highly variable and none were significantly different from 0. Nevertheless the higher Morisita index ratio compared to other indices seems to indicate a differentiation made by taxa abundance rather than by replacement or nestedness.

### Trait analysis

The traits studied tended to be similar in mound and control but different in pit (Table 3 and Figs 6 - 8). Considering alimentation traits (Fig 6), the proportion of collectors and scrapers was lower in the mound than in the pit and the proportion of filterers was lower in pit than either in mound or control. Predators were more present in mound than in pit and control, whereas shredders were less abundant in mound than in pit and control. However, food traits, related to alimentation aspects, did not highlighted significant differences. Substrate preference analysis (Fig 7) showed a similar pattern for litter and mud with higher proportion in pit than in mound. Preference for gravel and sand was more important in pit, followed by control and then mound. Finally, preference for slabs, blocks, stones and pebbles was highest in mound and lowest in pit. The smallest size category (Fig 8) was more represented inside the pit, followed by control and mound, whereas larger size categories followed the inverse pattern.

**Table 3.** Binomial mixed models results comparing frequency of selected traits between zones with nest identity as a random effect. With \**P* ≤ 0.05, \*\*\**P* ≤ 0.001.

|  | Zone | Df | Z value | P |
| --- | --- | --- | --- | --- |
| Food | Mound (reference: control) | 2 | -1.016 | 0.31 |
|  | Pit (reference: control) | 2 | -0.614 | 0.539 |
|  | Mound (reference: pit) | 2 | -0.401 | 0.688 |
| Alimentation | Mound (reference: control) | 2 | 1.877 | 6.05e-2 |
|  | Pit (reference: control) | 2 | -2.201 | 2.77e-2 * |
|  | Mound (reference: pit) | 2 | 4.063 | 4.84e-5 *** |
| Substrate | Mound (reference: control) | 2 | -6.918 | 4.57e-12 *** |
|  | Pit (reference: control) | 2 | 10.779 | <2e-16 *** |
|  | Mound (reference: pit) | 2 | -17.391 | <2e-16 *** |
| Size | Mound (reference: control) | 2 | 5.682 | 1.33e-08 *** |
|  | Pit (reference: control) | 2 | -11.63 | <2e-16 *** |
|  | Mound (reference: pit) | 2 | 16.71 | <2e-16 *** |

**Fig 6.**
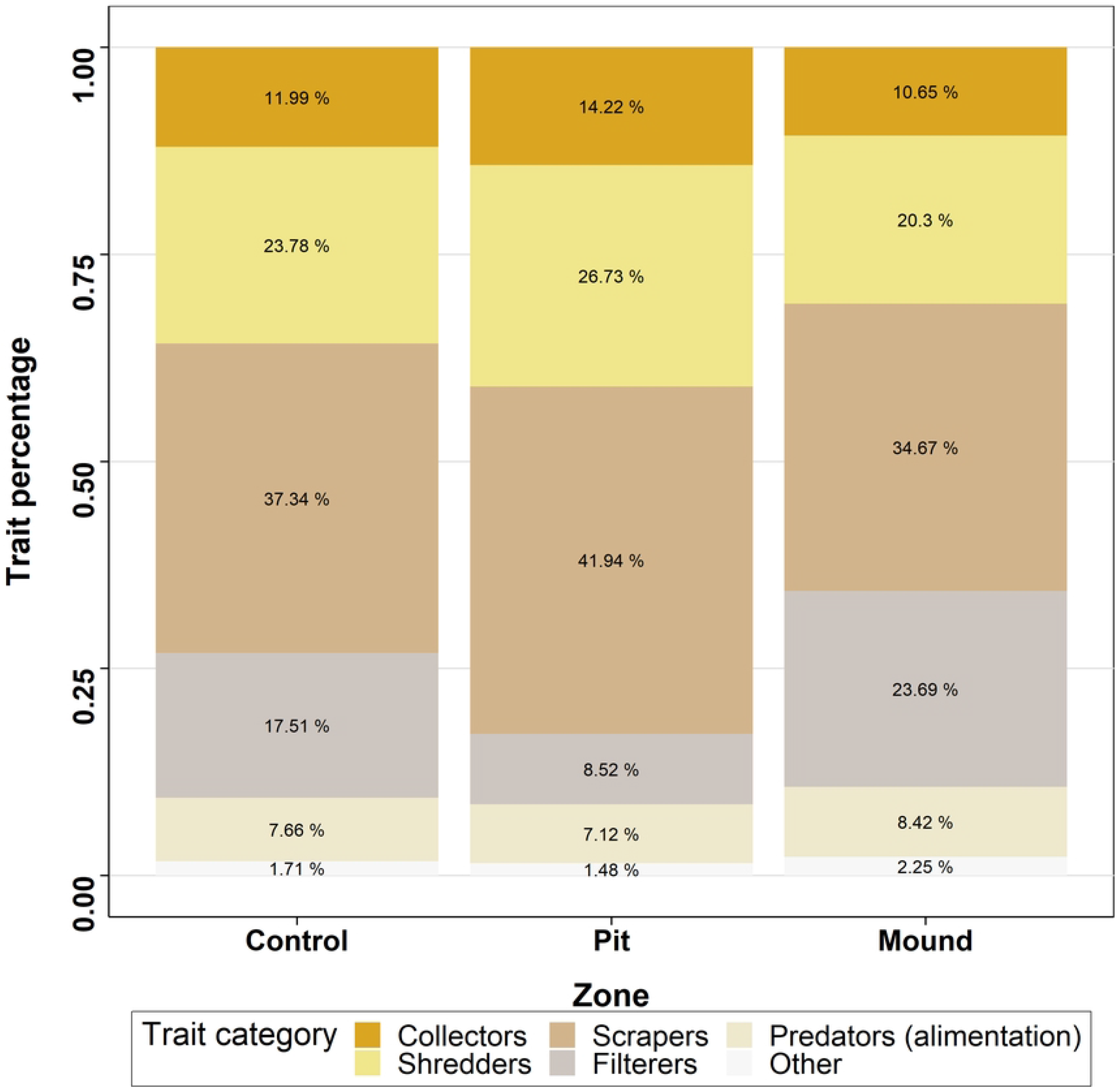
Median percentage of macroinvertebrate alimentation traits for the three zones studied.

**Fig 7.**
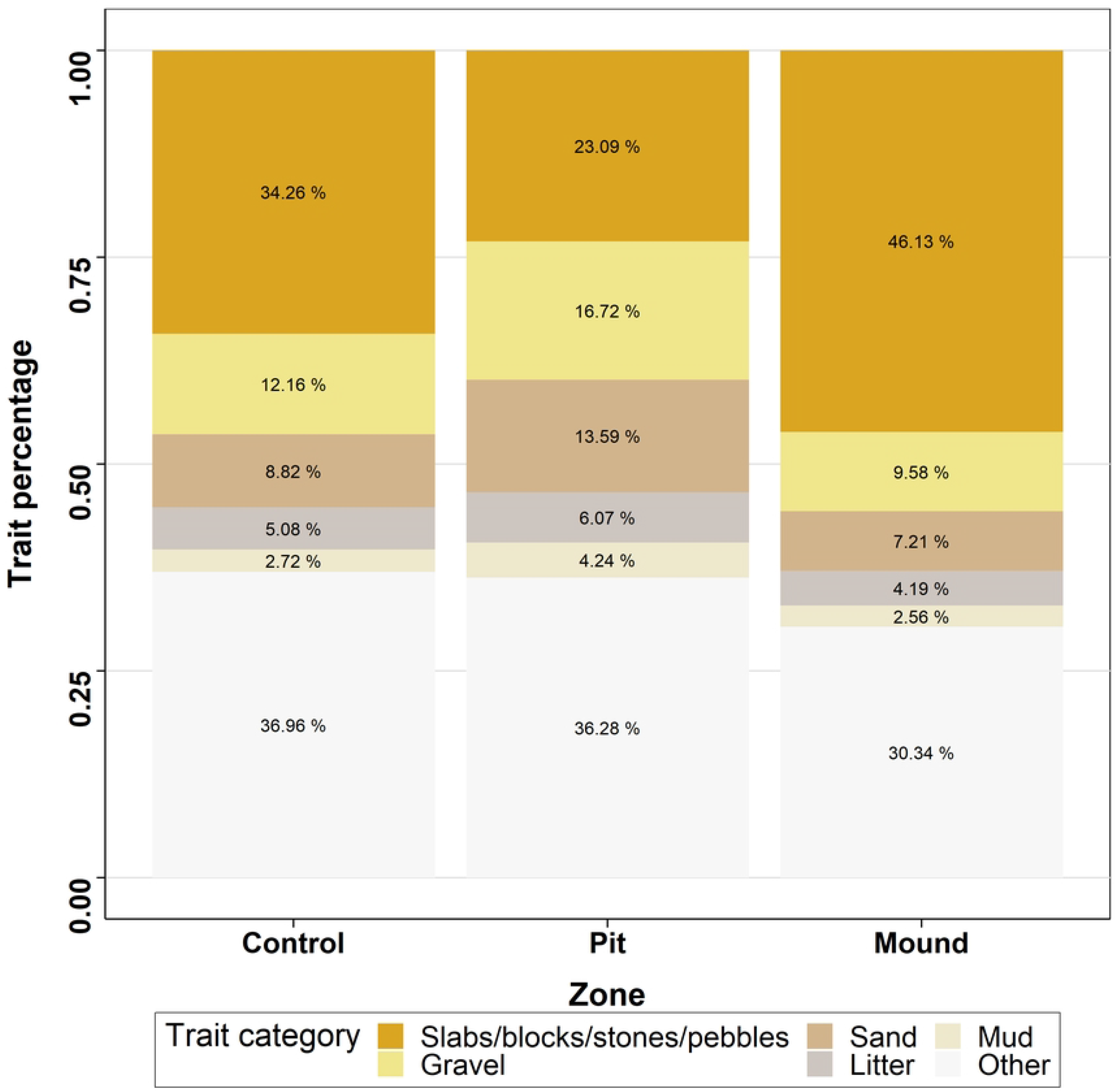
Median percentage of macroinvertebrate substrate preferences for the three zones studied.

**Fig 8.**
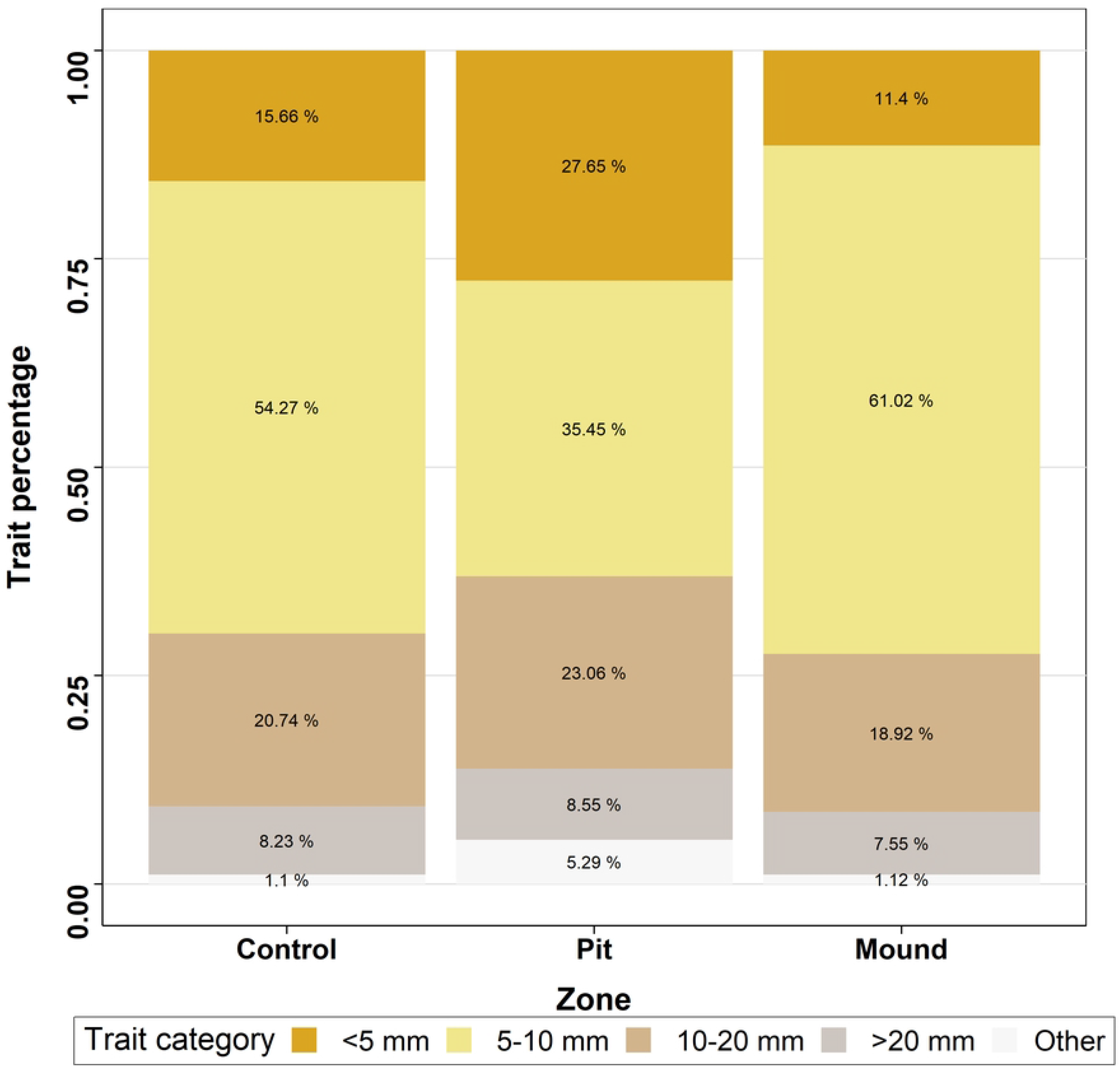
Median percentage of macroinvertebrate size traits for the three zones studied.

### Relationship between traits and nest characteristics

Redundancy analysis (Fig 9) yielded a significant model (*F* = 10.876, *Pr*(> *F*) = 0.001), although the only significant term was the current (*F* = 46.8357, *Pr*(> *F*) = 0.001), depth and depth difference being marginally significant (*F* = 2.4535, *Pr*(> *F*) = 0.077 and *F* = 2.6966, *Pr*(> *F*) = 0.077 respectively). The ellipses indicated a separation of pit from mound and control structured by current and “< 5 *mm*”, “*scrapers*”, “*sand*” and “*gravel*” trait categories belonging to the pit ellipse. Control and mound were not clearly separated but “*5-10 mm*”, “*slabs, blocks, stones and pebbles*” and “*filterers*” belong to the mound ellipse. Those results are consistent with the specific trait analyzes (Figs 6 - 8). Depth, if not the most discriminant variable, explained the separation of pit from other zones.

**Fig 9.**
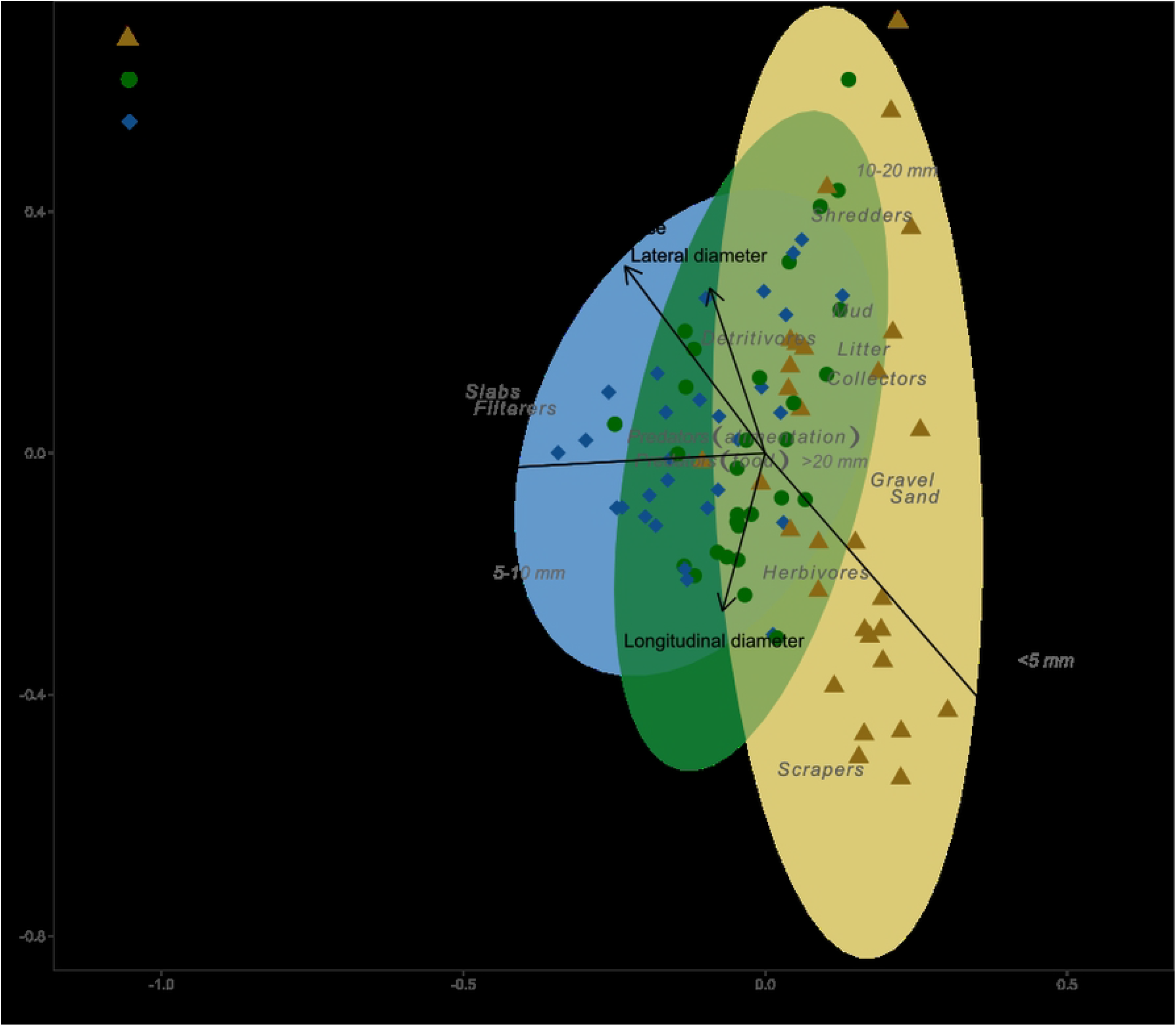
Redundancy analysis between nest characteristics and macroinvertebrate traits. Slabs = *slabs, blocks, stones and pebbles*.

Generalized Linear Models indicated that current affected neither the number of taxa nor the macroinvertebrate density (*tvalue* = 0.391; *Pr*(> |*t*|) = 0.697 and *tvalue* = 1.470; *Pr*(> |*t*|) = 0.1452 respectively). Depth influenced negatively the macroinvertebrate density (*tvalue* = −2.026; *Pr*(> |*t*|) = 0.0458) but not the number of taxa (*tvalue* = −0.821; *Pr*(> |*t*|) = 0.414).

## Discussion

Firstly, our results globally showed that habitat diversity and structural heterogeneity due to lamprey nesting activities modify several aspects of biodiversity. The clear distinction between nest zones regarding depth and riverbed current velocity implied biological heterogeneity, a non trivial result as previously highlighted [11]. The pit had a lower density of invertebrates and a lower number of taxa than the other zones, which did not differ between them, although the number of taxa tended to be higher at the control. Log response ratio of *α* Shannon index showed a similar trend concerning the diversity inside the pit. Those results indicate that nest building reduced the local abundance and diversity of macroinvertebrates in the pit, at least during the first weeks after nest digging, which are the most significant from the point of view of the lamprey, as larvae abandon the nest shortly after hatching [36]. Patches of fine substrate such as the pit tend to be less diverse than zones with coarser grain size [37]. Reduced invertebrate abundance and richness have been also reported in nests of the largemouth bass (*Micropterus salmoides*) [38] and the pink salmon (*Oncorhynchus gorbuscha*) [39] during the spawning season, which was attributed to spawning-related disturbance. On the other hand, Hogg et al. [22] reported that the density of invertebrates in sea lamprey nest mounds were twice that found in pits and 77% higher than in control zone. Their results of reduced macroinvertebrate pit abundance are consistent with ours but we did not observe an increased mound density. An explanation for the important density in mounds could be the dominance of the Chironomidae within the river studied, not found for the Nive River (Fig 10 in supporting information). The difference between our results and theirs seems not to derive from differences in macroinvertebrate assemblages, as these were relatively similar between both studies. A significantly higher number of taxa in control than in mound and pit indicates that sea lamprey nests create heterogeneity but do not increase local species diversity.

In a local scale, i.e. the zone, we found for each pit and mound several taxa not present in the control (see Fig 11 in supporting information). Indeed, whereas control is more complex and offers more substrate types with its unsorted grain size - likely to provide suitable habitat for more taxa - [40], mound and pit provide larger areas of some specific characteristics, including substrate. This wider area may potentially allow the establishment of more taxa sharing similar ecological preferences, when these would be less represented and therefore potentially absent or very rare in the control. Considering the nest scale, rare taxa should be found more easily, either in mound or in pit considering their preferences than in control (considering a same sample size). At a larger scale, occurrence of rare species and so species diversity would be higher in zones with nests than in zones without them.

In addition to increasing the structural and biological heterogeneity, sea lamprey nests seem to shape invertebrate assemblages. Indeed, three traits over the four studied showed differences between nest and the control zones, although the direction of change did not always follow our expectations. In particular, we expected herbivores to increase in mounds, as epilithic algae tend to be favored by shallow water, coarse sediments and moderately fast flow [41, 42], but no significant difference was found globally for alimentation traits, perhaps because of lack of time for algae to grow on overturned cobbles. The increased proportion of collectors in pits was expected, as these are a preferential place for organic matter deposits. Similarly, collectors tend to be more abundant in fine substrate, which collected more detritus [43]. The higher concentration of organic matter might also explain the higher abundance of shredders and scrapers in pit than in mound. Also, according to our expectations, filterers were more represented in the mound, as they are favoured by fast-flowing areas [44], particularly during low flow conditions [45]. Those results reflect the different dynamics of food provided within the nest zones.

Not surprisingly given the differences in substrate among zones, invertebrates also differed in their preferred substrate traits. Macroinvertebrates associated with coarse substrate were especially abundant in mounds, whereas taxa associated with fine substrate were more abundant in pits. It must be noted that sand was almost absent in the study reach apart from lamprey nests and some marginal areas, which suggests that nesting lampreys create patches of habitat for lentophylic species in river stretches where these species could not dwell otherwise.

The macroinvertebrate size traits showed a greater proportion of small invertebrates (< 5 mm) within the pit, including *Esolus sp*., the most abundant taxon in pit (Fig 10 in supporting information) and *Psychomyia pusilla*. This result could be explained by the carrying capacity of fine substrate. A fine substrate has less shelters than a coarser one. Shelters of fine substrate are more suitable for small size range of macroinvertebrates, whereas bigger macroinvertebrates are more prone to find shelter in the coarser substrate of the mound or the control. Bêche et al. [46] suggested a better exploitation of refugia for small sized macroinvertebrates. Furthermore, small body size could be a possible resistance trait to disturbances such as riverbed movement [47].

Our results demonstrate that sea lamprey creates physical heterogeneity, which then enhances biological heterogeneity both in assemblage composition and function. The heterogeneity demonstrated in this study was created at the nest scale, but local-scale processes can have a major impact at higher scales such as the reach scale [48]. In the case of lamprey nests in the Nive River, pits average 1.15 *m*^2^ and so the total modified streambed area averages 34.5 *m*^2^, representing 4% of the river streambed in our studied zone. With an average reduction of 30% of macroinvertebrate density in pit (compared to control) a total reduction of 1.2% of density is expected in the studied zone. In a river reacting in a similar way to the Nive, this surface with decreased macroinvertebrate density does not seem to be compensated by a higher density in the mound and so, sea lamprey nests globally decrease the macroinvertebrate density.

Measuring the effect of lamprey nests on reach-scale macroinvertebrate density would require a BACI design [49] including periods and reaches with and without nests. If invertebrates affected by lamprey nesting responded with small-scale movements, the result would be an increase in macroinvertebrate density in control samples of reaches with nests compared to control samples of reaches without nests. If the macroinvertebrates responded drifting, the results would be more complex to interpret and depend on the distribution of nests of sea lamprey. An increase of macroinvertebrate densities could be observed if spawning sites occur upstream. Macroinvertebrates drifting from those nests could colonize and increase the density of downstream control sites. When a spawning ground is located on the upper limit of the area colonized by lampreys, drift will not affect the adjacent area of this spawning ground but may affect downstream sites.

Our results highlight the need for a better understanding of the effects of nest-building species on the river ecosystem. Unlike salmon, whose effects are relatively well-studied [50, 51, 52], the consequences of sea lamprey spawning are still poorly known. Whereas lamprey nests only represent 4% of the river streambed in our study, the occupied area may be much higher, especially for invasive populations of the Great Lakes [53] or below impassable dams, likely to dramatically increase the number of spawners in immediately downstream spawning grounds [24, 54]. Studies exist on the effects of carcasses [55, 23] or substrate modification [22] on a local scale, but not at the river or at the global scales. However, macroinvertebrates are at the bottom of the food chain [56]. Such basal position implies an important relationship with ecosystem components. Firstly, macroinvertebrate assemblages can have an important influence on numerous processes [56], such as nutrient cycles [57, 58, 59], primary productivity [60], decomposition [61, 62, 63], and translocation of materials [64, 65]. Then, they are an important resource for organisms belonging to higher trophic levels such as fish species. Macroinvertebrates are considered as the most important source of food, widespread in all freshwater ecosystems [66].

In our study some trait aspects can provide clues about the effects of sea lamprey on general ecosystem functioning in the river. Increased proportion of shredders in the pit suggests they find suitable conditions such as increased plant debris. On a more global scale a certain amount of material (depending on the nest density and count) could be retained by nests in spite of being carried away by the current, and then consumed by macroinvertebrates. Seeing this impact in late spring when litter input is reduced also indicates that the proportion of shredders should be more important in autumn, where these inputs can represent 73% of annual allochtonous inputs (in temperate deciduous forests; Abelho and Graça [67]). But this retention can occur only if the lamprey nests remain in autumn, without being filled by floods. Changes in streambed complexity created by nests persist during the autumn [22] but a change in hydrological conditions is likely to affect this process. Filterers are another trait category possibly affecting organic matter processing, being one pathway through which carbon and nutrients are transferred from the water column to the sediments [68]. Filterers being more present in proportion in the mound than in the control suggests that mounds are hotspots of nutrient processing. Therefore, sea lamprey nests could have an important effect on nutrient spiraling in streams, but the persistence of their influence on ecosystem functioning until the next spawning season remains to be assessed. Although nests may persist considering their physical structure, it is likely that the subsidies provided by eggs affect macroinvertebrate colonization dynamics and the distribution of functional groups, as predation was depicted in other species [69, 70]. An assessment of the effects of eggs deposited in the pit on functionality should be done to determine what is their potential contribution to the observed differences.

All these possible different effects imply that the modifications of macroinvertebrate assemblages by sea lampreys highlighted here and in previous studies [22, 23] are likely to modify the general functioning of the rivers. However, it is difficult to determine these consequences precisely due to the complexity inside macroinvertebrate assemblages. Such assessment requires specific studies on the components previously described. We advocate for studies focusing on these effects, important in a context of endangered native populations, likely to disappear in some rivers and consequently modifying their dynamics. In addition, negative impacts of invasive populations through population reduction of their hosts may be mitigated by a positive effect through invertebrate diversity.

## Acknowledgements

Functioning was financed by Pôle Gestion des Migrateurs Amphihalins dans leur Environnement. M.D. PhDs was financed by Univ. Pau and Pays Adour and UPV/EHU. Field work used resources from the IE ECP Experimental Facility of the UMR Ecobiop [71].

## Author contributions

Conceptualisation: MD, CT and AE; Developing methods: MD, CT, JR and AE; Conducting the research: MD, CT, JR, AE, MP; Data analysis and interpretation: MD helped by AL, AE, CT; Preparation of figures and tables: MD; Writing: MD, critically revised by CT, AL and AE.

## Data availability

Data and code to run the analysis are available in the INRAE dataverse repository at https://doi.org/10.15454/BEHOQK, under the name “Sea lamprey nests promote the diversity of benthic macroinvertebrate assemblages”

## Supporting information

Fig 10. Median of the abundances for the 10 most abundant taxa within each zone (Control, Pit, Mound).

Fig 11. Number of exclusive taxa in mound or pit compared to the control.

With \*\**P* ≤ 0.01.

